# Fibrillarin modulates fetal hemoglobin silencing

**DOI:** 10.1101/2024.06.25.600532

**Authors:** Dongliang Wu, Qixiang Li, Sipei Qiu, Chan Guo, Feng Li, Wenbing Shangguan, Wenyang Li, Dongjun Yang, Xingjun Meng, Mengying Xing, Bing Chen, Lingdong Kong, David C. S. Huang, Quan Zhao

## Abstract

Decoding the molecular mechanisms underlying human fetal (γ) globin gene silencing impacts therapeutic strategies for β-thalassemia and sickle cell disease. Here, we identified a nucleolar protein, fibrillarin (FBL), which mediates the methylation of glutamine104 in histone H2A and functions as a repressor of the γ-globin gene in cultured erythroid cells, including those from β-thalassemia patients. Conditional *Fbl* depletion in adult β-YAC transgenic mice or in βIVS-2-654-thalassemic mice reactivated the human γ-globin gene or murine embryonic globin expression, respectively, which corrects hematologic and pathologic defects in β-thalassemic mice. We showed that FBL plays a dual role in activating *BCL11A* expression and repressing γ-globin gene expression, which is dependent on its histone methyltransferase activity. Our study may provide an alternative strategy for therapeutic targeted treatment of β-hemoglobinopathies.

## Introduction

It has been a long-term goal to define the mechanism underlying the switch from fetal (HbF, α2γ2) to adult hemoglobin (HbA, α2β2), driven by the desire to reverse this switch as a means to treat β-thalassemia and sickle cell disease (SCD) (*1–3*). This elegant work includes discoveries of the transcription factors BCL11A, LRF (ZBTB7A), NFI, and other cofactors that participate in regulating the expression of the fetal globin genes *HBG1/2* (*4–15*). These studies, conducted over decades, have culminated, in 2023, in the discovery of Casgevy, the first ever FDA-approved gene therapy product targeting the BCL11A enhancer for the treatment of β-thalassemia and SCD (*16–18*). However, despite therapeutic success, currently, the necessary infrastructure, resources, and cost may render gene therapy approaches largely inaccessible.Alternative strategies and more feasible treatments for β-hemoglobinopathies are highly needed.

## Results

### Identification of FBL as a γ-globin repressor

In our previous studies, we showed that the arginine methyltransferase PRMT5 interacted with the transcription factor LYAR to repress human γ-globin expression (*19–22*). To further identify other potential proteins that interact with LYAR, we initially immunoprecipitated LYAR from the cellular extracts of K562 cells overexpressing Flag-tagged LYAR, and LYAR-associated proteins were analyzed by mass spectrometry. As expected, the PRMT5-partner proteins Nucleolin and MEP50 (*23, 24*) were identified (Fig. S1A). In addition, Fibrillarin (FBL), a conserved nucleolar protein with methyltransferase activity (*25, 26*), was identified as a novel LYAR-interacting protein (Fig. S1A). We then confirmed this interaction in both K562 and HUDEP-2 cells by coimmunoprecipitation (Co-IP) experiments (Fig. S1B and C). Furthermore, GST pull-down assays showed that FBL interacted with LYAR directly (Fig. S1D). Since LYAR has previously been shown to bind the γ-globin gene promoter, we tested whether FBL bound to the γ-globin promoter. We performed ChIP-reChIP experiments and found that FBL and LYAR bound to the γ-globin promoter in both K562 cells and HUDEP-2 cells (Fig. S1E and F). Moreover, LYAR knockdown disrupted FBL binding to the γ-globin promoter without affecting FBL expression (Fig. S1G, H and I). Since FBL harbors methyltransferase activity, which specifically triggers the methylation of glutamine 104 in histone H2A (H2AQ104me) (*25*), we correspondingly observed that the enrichment of H2AQ104 at the γ-globin promoter was significantly reduced in LYAR-deficient cells compared to that in SCR control cells (Fig. S1G and H). Together, these data indicate that FBL interacts with LYAR in erythroid cells and that the binding of FBL to the γ-globin promoter is LYAR dependent.

To examine the impact of FBL on γ-globin gene (*HBG*) expression, we first generated two stable FBL-knockdown HUDEP-2 cell lines (FBL-KD1 and FBL-KD2) with specific shRNAs (Fig. 1A). We found that knockdown of FBL significantly increased *HBG* mRNA levels, γ-globin protein expression, and the percentage of HbF-expressing HUDEP-2 cells compared to scrambled control (SCR) cells (Fig. 1B to D and Fig. S2A). As expected, the global H2AQ104me level was significantly lower in FBL-knockdown HUDEP-2 cells than in SCR cells (Fig. 1C). As such, we observed coordinated significant reductions in FBL enrichment, H2AQ104me enrichment, but increase in RNA polymerase II (RNA Pol II) occupancy on the γ-globin promoter in FBL-knockdown HUDEP-2 cells compared to SCR cells (Fig. S1J and K). Notably, *HBB* mRNA levels did not substantially change (Fig. S2A). In fact, FBL depletion did not affect the erythroid maturation process, as shown by cell surface phenotyping (CD 235a and CD71) and Wright‒Giemsa staining at the end of differentiation (Fig. S2B and C).

**Fig. 1.**
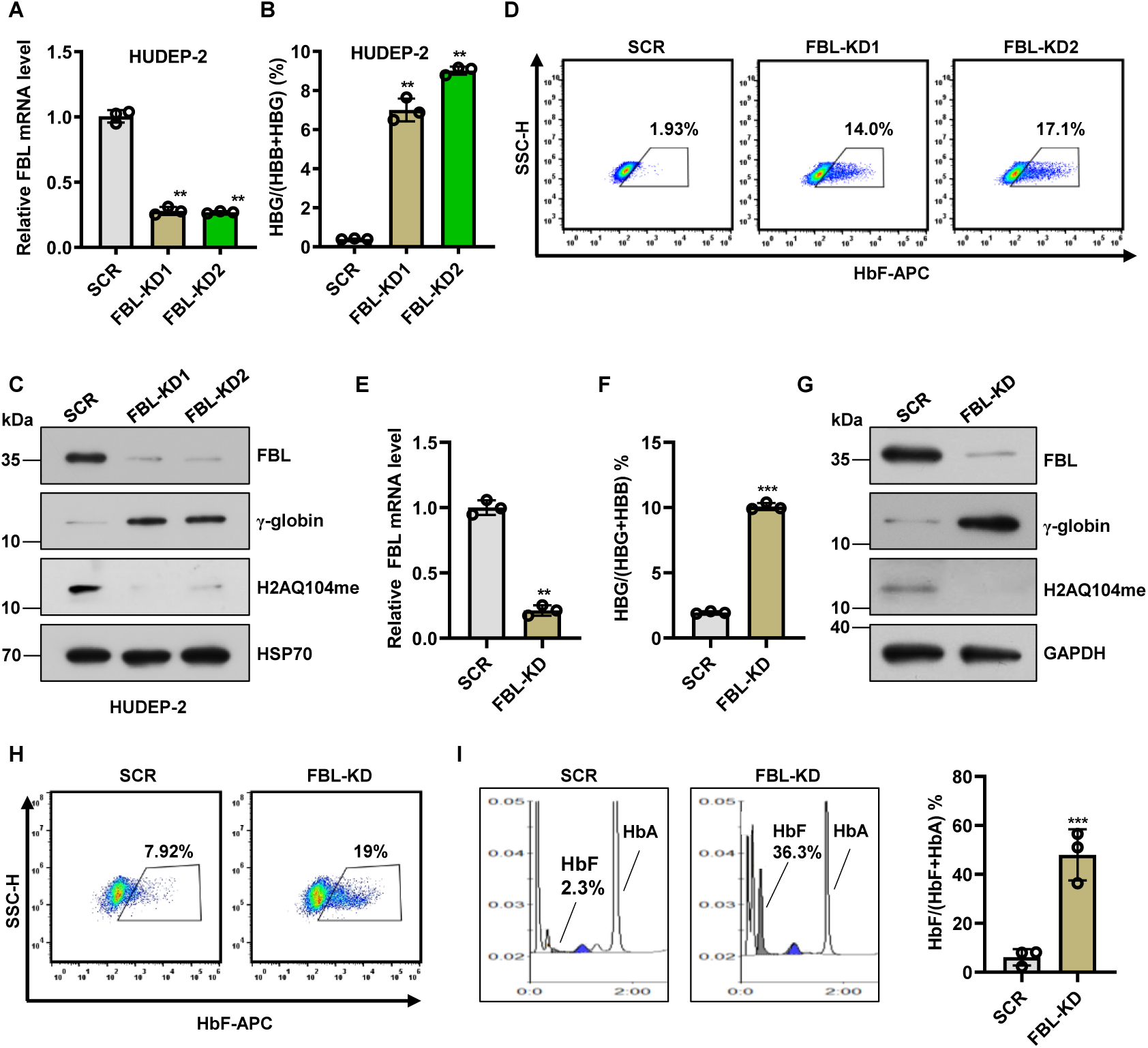
FBL disruption induces γ-globin expression in HUDEP-2 cells and primary adult erythroblasts. **(A)** The relative FBL mRNA expression in FBL-KD1, FBL-KD2, or SCR HUDEP-2 cells was analyzed by real-time qPCR. The results were normalized to GAPDH mRNA and are shown as the mean ± SD from three independent experiments. \*\**P* < 0.01. **(B)** Real-time qPCR was used to determine the HBG/(HBG + HBB) ratio in the indicated HUDEP-2 cells. The results were normalized to GAPDH mRNA and are shown as the mean ± SD from three independent experiments. \*\**P* < 0.01. **(C)** Western blot analysis of FBL, H2AQ104me proteins, and γ-globin in the indicated cells. HSP70 served as a loading control. **(D)** The percentage of HbF-immunostained F-cells was detected by flow cytometry. **(E)** The relative FBL mRNA expression in FBL-KD or SCR CD34^+^ HSPCs was analyzed by real-time qPCR. The results were normalized to GAPDH mRNA and are shown as the mean ± SD from three independent experiments. \*\**P* < 0.01. **(F)** Real-time qPCR was used to determine the HBG/(HBG + HBB) ratio in the indicated HSPCs. The results were normalized to GAPDH mRNA and are shown as the mean ± SD from three independent experiments. \*\*\**P* < 0.001. **(G)** Western blot analysis of FBL, H2AQ104me, and γ-globin in the indicated HSPCs. HSP70 served as a loading control. **(H)** The percentage of HbF-immunostained F-cells was detected by flow cytometry. **(I)** Fetal hemoglobin (HbF) and adult hemoglobin (HbA) levels were analyzed by HPLC in SCR or FBL-KD HSPCs (left). The proportion of HbF relative to adult globin (HbA) was quantitated on day 18 (right). The results are shown as the mean ± SD from three independent experiments. \*\*\**P* < 0.001.

### FBL disruption induces γ-globin expression in primary human erythroid progenitor cells from healthy donors and β-thalassemia patients

To further validate the above results, we assessed the effect of FBL on γ-globin expression in CD34^+^ adult hematopoietic stem and progenitor cells (HSPCs) (Fig. S3A). We constructed stable FBL-knockdown (FBL-KD) CD34+ cell lines with a specific lentiviral shRNA (Fig. 1E and G). We showed that FBL silencing significantly increased the ratio of *HBG/(HBG+HBB)* mRNA from 1.9% to 10% but did not alter *HBB* mRNA levels (Fig. 1F and Fig. S3B). Consistently, FBL silencing significantly increased the γ-globin protein, the percentage of HbF-expressing cells from 7.9 to 19%, and HbF levels from 2.3% to 36.3%, as assessed by high-performance liquid chromatography (HPLC) (Fig. 1G, H, I and Fig. S3B). Notably, no changes in cell viability, expression of maturation markers, or cell morphology were observed in FBL-depleted cells compared to SCR cells (Fig. S3C, D, and E). To examine the role of FBL in the developmental silencing of γ-globin genes, we determined FBL expression levels in cord blood and adult HSPCs. We found that both the messenger RNA and protein levels of FBL were significantly greater in adult HSPCs (in the γ-globin ‘off’ state) than in cord blood cells (in the γ-globin ‘on’ state) (Fig. S3F and G). We further examined FBL protein levels during the maturation of primary erythroblasts and found that FBL protein was constitutively expressed at a high level from early stages to late mature erythroblasts and markedly increased in more mature (days 11–15) erythroblasts (Fig. S3H). Together, these results indicate that FBL repressed human γ-globin in HSPCs.

To explore the function of FBL in CD34^+^ HSPCs from β-thalassemia patients with IVS-2-654 (C>T) mutations (*27, 28*), we disrupted FBL and examined the expression of γ-globin. Consistent with our findings in HSPCs from healthy donors, disruption of FBL increased γ-globin gene expression, γ-globin protein expression, and the percentage of HbF-expressing cells to levels similar to those following disruption of BCL11A (Fig. S4A, B and C). We detected decreased levels of H2AQ104me and decreased levels of BCL11A upon FBL knockdown (Fig. S4B). In addition, we found that the size of erythroid cells from β-thalassemia patients increased after differentiation upon FBL depletion (Fig. S4D). Taken together, these results indicate that FBL disruption induces γ-globin expression in primary human erythroid progenitor cells from healthy donors and β-thalassemia patients.

### Fbl depletion elevates human γ-globin and mouse embryonic globin gene expression in adult β-YAC mice

Mouse and human β-globin-like loci are highly conserved and share most functional elements (*15, 29, 30*). To test the impact of FBL on globin gene expression during normal development in vivo, we constructed a mouse line carrying an *Fbl* floxed allele (with a loxP site flanking exons 2 to 4, Fig. S5A) and an interferon-inducible Mx1-Cre allele (*31*) and carrying a yeast artificial chromosome transgene harboring the human β-globin gene cluster (β-YAC) (*32*) (Fig. 2A). This mouse model permits examination of the contribution of Fbl to the silencing of the human g-globin gene in adult mice upon hematopoietic conditional loss of *Fbl*. Mice (6–8 weeks old) were subjected to 4 courses of poly (I:C) treatment (Fig. 2A). At the end of treatment followed by recovery for one week, the efficient excision of exons 2 to 4 in mouse bone marrow was verified by PCR and compared with control mice (Fig. 2B). Western blot analysis also confirmed that Fbl was absent from the bone marrow of *Fbl*^fl/fl^::Mx1-*Cre*^+^::β-YAC mice (Fig. 2C). These results indicated that deletion of *Fbl* exons 2 to 4 abolished the expression of the Fbl protein. No overt changes in blood counts were observed after depletion of Fbl (Fig. S5B). Importantly, in *Fbl*^fl/fl^::Mx1-*Cre*^+^::β-YAC mice, *HBG* expression was significantly increased, but there was no significant change in HBB expression (Fig. 2D and Fig. S5C). Similarly, we also observed the induction of mouse embryonic εy-globin and βh1-globin genes but no changes in mouse adult β-globins (βmajor and βminor) (Fig. 2E and Fig. S5D). Notably, *Fbl* depletion did not affect erythroid maturation, as determined by the detection of CD71 and Ter119 markers (Fig. S5E). Together, these results show that Fbl modulates mouse embryonic globin gene silencing, consistent with the function of FBL in regulating the human γ-globin gene.

**Fig. 2.**
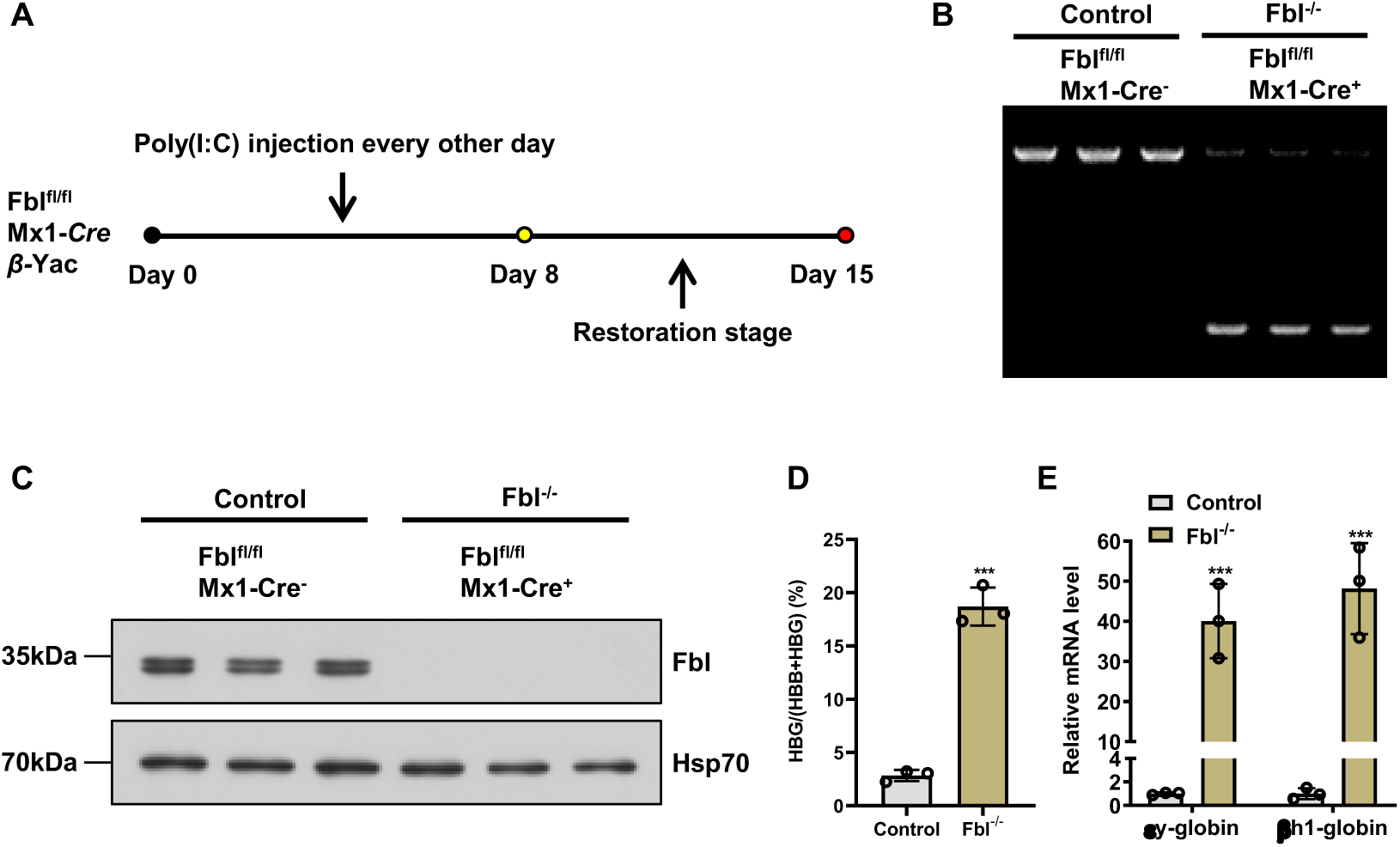
Fbl depletion elevates human γ-globin and mouse embryonic globin gene expression in adult β-YAC mice. **(A)** Experimental scheme for inducing Fbl knockout in adult *Fbl*^fl/fl^::Mx1-*Cre*^+^::β-YAC mice by interferon-mediated deletion induced by an 8-day course of poly (I:C) intraperitoneal (i.p.) injections (arrows). **(B)** PCR analysis showing the deletion efficiency in bone marrow cells from control (*Fbl*^fl/fl^::Mx1-*Cre*^-^::β-YAC) and *Fbl*^-/-^ knockout (*Fbl*^fl/fl^::Mx1-*Cre*^+^::β-YAC) mice after induction. An ethidium bromide-stained agarose gel is shown. **(C)** Western blot of Fbl protein in bone marrow cells from control and *Fbl*^-/-^ knockout mice after induction. HSP70 served as a loading control. **(D)** Real-time qPCR was used to determine the HBG/(HBG + HBB) ratio in the indicated mice. (**E**) Expression of mouse embryonic εy-globin and βh1-globin mRNA in the indicated mice. The results were normalized to β-actin mRNA and are shown as the mean ± SD from three independent experiments. \*\*\**P* < 0.001.

### Depletion of Fbl corrects anemia in a mouse model of beta-thalassemia

To further explore the role of Fbl in a β-thalassemia mouse model, we used a disease model in which the two normal adult murine β-globin genes were replaced by a single copy of the human βIVS-2-654 mutant gene, which has a moderate to severe form of β-thalassemia (*33, 34*). We generated an adeno-associated virus (AAV) carrying Fbl-specific shRNA to disrupt Fbl in mouse bone marrow. βIVS-2-654 β-thalassemia mice (8 weeks old) received intramedullary virus injection. To evaluate the efficiency of AAV carrying Fbl-specific shRNA in bone marrow, we performed western blot analysis and showed that Fbl expression in the bone marrow of β-thalassemia mice in the AAV-Fbl KD group was largely silenced compared to that in the AAV-SCR or wild-type groups (Fig. 3A). RBC morphology was significantly improved in the AAV-Fbl-KD group but not in the AAV-SCR group (Fig. 3B). Complete blood analysis of blood samples indicated that anemia was significantly improved based on blood hemoglobin levels in the AAV-Fbl-KD mice but not in the AAV-SCR mice (Fig. 3C). We also found that reticulocyte counts in AAV-Fbl-KD mice were similar to those in wild-type mice but markedly increased in the AAV-SCR group (Fig. 3D). Consistent with our findings in adult β-YAC mice, mouse embryonic εy-globin and βh1-globin gene expression was markedly increased in the AAV-Fbl-KD group (Fig. 3E), whereas mouse adult βmajor-globin and βminor-globin gene expression was slightly reduced (Fig. S6A). In addition, we also found significantly less splenomegaly in AAV-Fbl-KD mice than in AAV-SCR mice (Fig. 3F, G, and Fig. S6B). Due to extramedullary hematopoiesis, the red pulp of the spleen expanded (leading to splenomegaly), and the amount of white pulp decreased in the AAV-SCR model mice, which was reversed by depletion of Fbl in the AAV-Fbl-KD group (Fig. 3H). The presence of Fbl depletion-alleviated splenomegaly and a corrected spleen structure in β-thalassemia mice are consistent with the blood indices. These results indicate that *Fbl* depletion is able to correct anemia in a mouse model of beta-thalassemia.

**Fig. 3.**
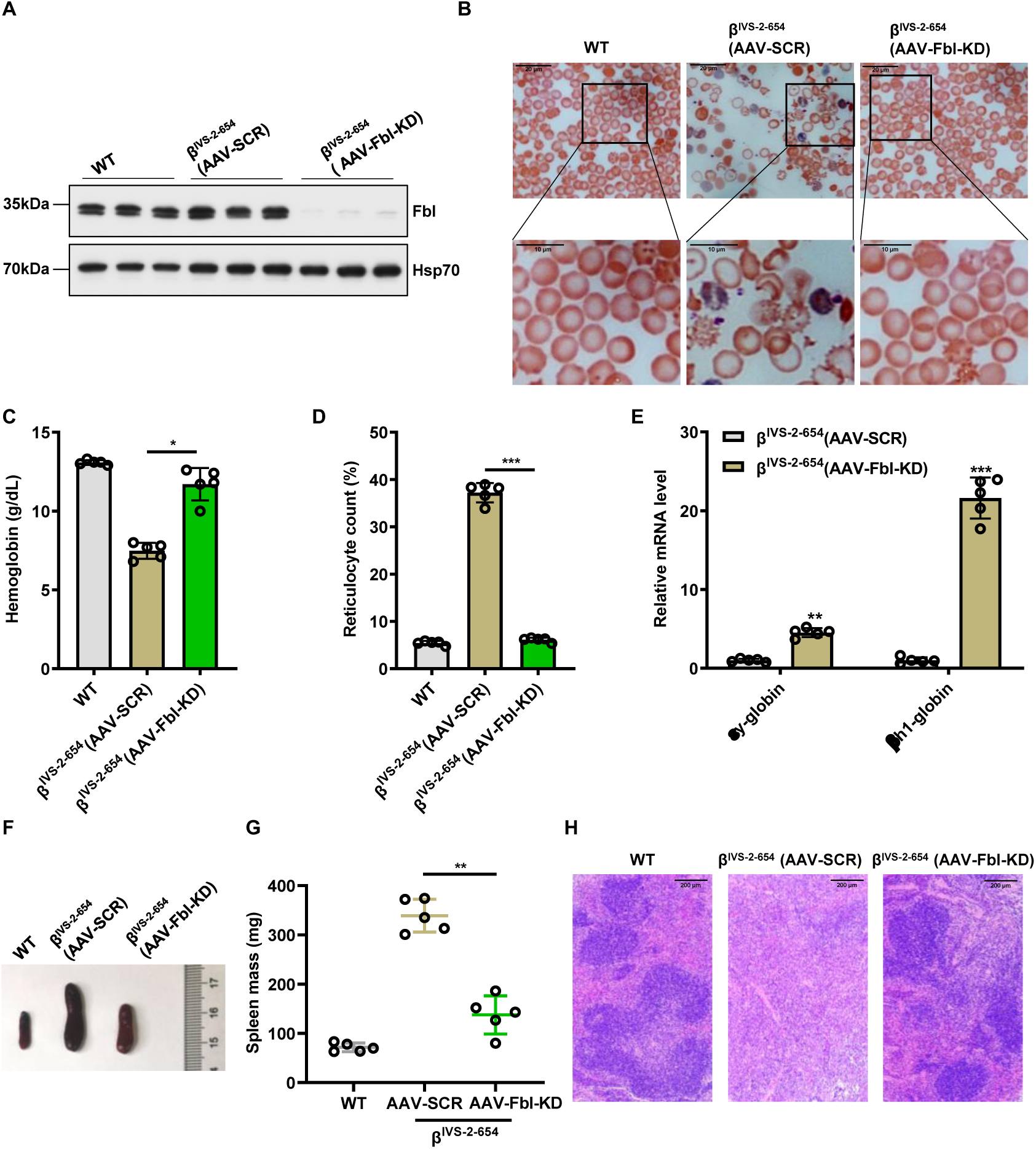
Depletion of Fbl corrects anemia in a mouse model of beta-thalassemia. **(A)** Western blot analysis of Fbl bone marrow cells from wild-type (WT), AAV-SCR-treated, or AAV-FBL-KD-treated thalassemic mice. HSP70 served as a loading control. **(B)** Blood smears were used to analyze RBC morphology in WT, AAV-SCR-treated or AAV-FBL-KD-treated thalassemic mice. Scale bar, 10 μm. **(C and D)** Blood hemoglobin levels (**C**) and reticulocyte counts (**D**) were analyzed in WT, AAV-SCR-treated, and AAV-FBL-KD-treated thalassemic mice (n=5 per group). The results are shown as the mean ± SD. \**P* < 0.05; \*\*\**P* < 0.001. **(E)** Real-time qPCR showed the expression of mouse embryonic εy-globin and βh1-globin mRNA in AAV-SCR-treated or AAV-FBL-KD-treated thalassemic mice. The results were normalized to β-actin mRNA and are shown as the mean ± SD from five independent experiments. \**P* < 0.05; \*\**P* < 0.01; \*\*\**P* < 0.001. **(F and G)** Spleen weights (**F**) and spleen images (**G**) of WT, AAV-SCR-treated, and AAV-FBL-KD-treated thalassemic mice (n=5 per group). \*\**P* < 0.01. Additional images of the spleen are presented in Fig. S6. **(H)** Histopathology of spleen sections from WT, AAV-SCR-treated, and AAV-FBL-KD-treated thalassemic mice. Scale bar, 200 μm.

### FBL promotes BCL11A expression in erythroid cells

To determine whether FBL selectively affects the γ-globin regulatory circuitry, we performed RNA sequencing (RNA-seq) in FBL-knockdown or scrambled (SCR) control HUDPE-2 cells. We conducted differential expression analysis and found that the HBG1/2 genes were among the most upregulated globin genes in FBL-depleted cells (Fig. 4A). Notably, the expression of the embryonic-type α-globin gene HBZ was also upregulated when FBL was depleted (Fig. 4A). To our surprise, transcriptional repressors that are highly expressed in the adult stage, such as BCL11A, were markedly downregulated (Fig. 4A). Gene set enrichment analysis (GSEA) revealed that beta-thalassemia-related genes were significantly enriched in FBL-knockdown HUDEP-2 cells compared with SCR controls (Fig. 4B). These results strongly suggest an important role for FBL in globin gene regulation.

**Fig. 4.**
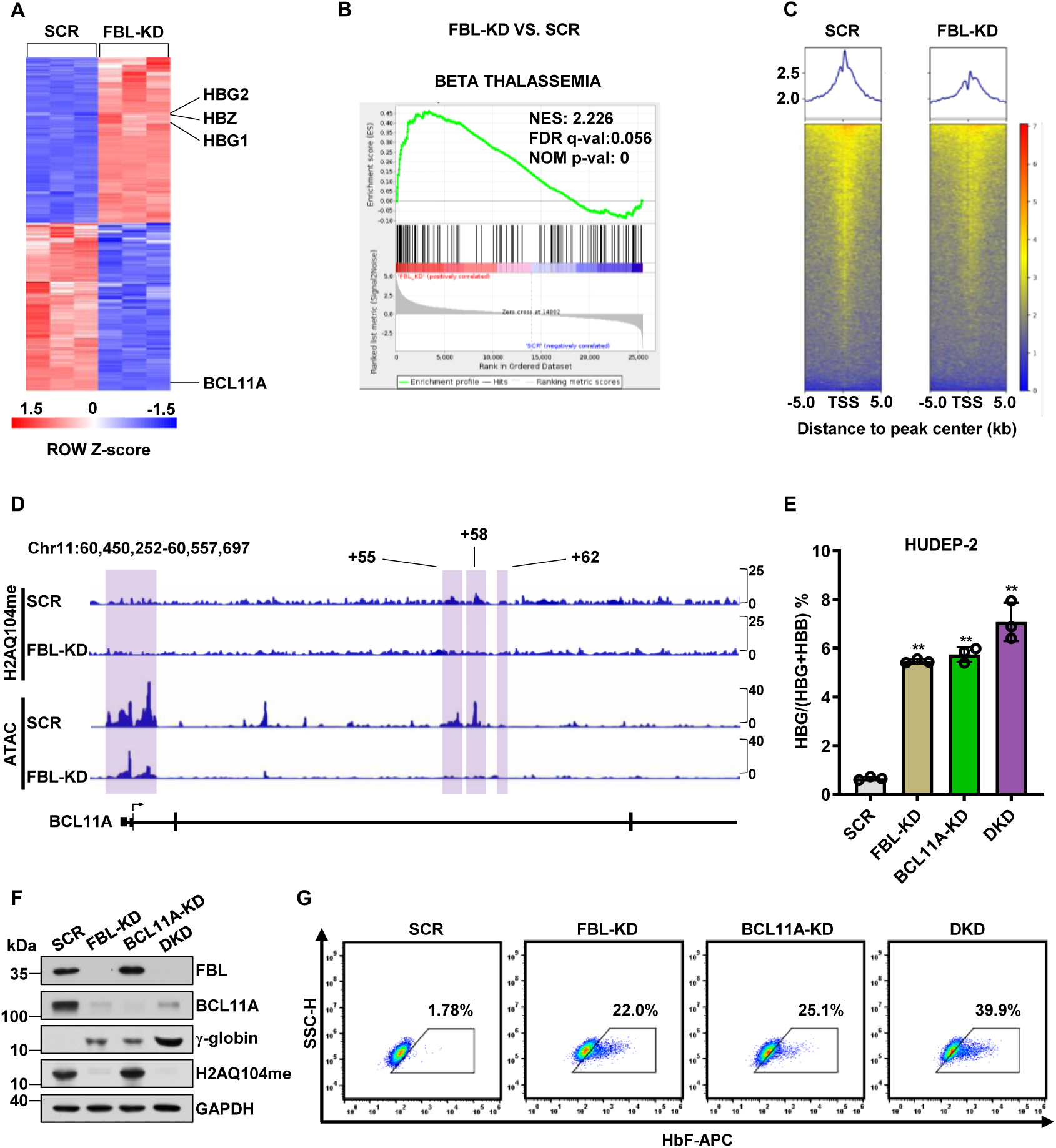
FBL promotes BCL11A expression in erythroid cells. (**A**) Hierarchical cluster analysis of all genes exhibiting differential expression in FBL knockdown (FBL-KD) or Scr control (SCR) HUDEP-2 cells. (**B**) Gene set enrichment analysis (GSEA) for beta-thalassemia in FBL-KD or SCR HUDEP-2 cells; n = 3 per group. (**C**) Visualization of FBL CUT&RUN peaks in SCR or FBL-KD HUDEP-2 cells by profile and heatmap plots. (**D**) Enrichment of H2AQ104me at the BCL11A locus in FBL-KD or SCR HUDEP-2 cells, as determined by CUT&Tag and analysis of open chromatin regions via ATAC-seq. **(E)** Real-time qPCR was used to determine the ratio of HBG/(HBG + HBB) in SCR, FBL-KD, BCL11A-KD, and double knockdown (DKD) HUDEP-2 cells. The results were normalized to GAPDH mRNA and are shown as the mean ± SD from three independent experiments. \*\**P* < 0.01. **(F)** Western blot analysis of FBL, BCL11A, γ-globin, and H2AQ104me in the indicated HUDEP-2 cells. GAPDH served as a loading control. **(G)** The percentage of HbF-immunostained F-cells in the indicated HUDEP-2 cells was determined by flow cytometry.

To probe the genomic occupancy of FBL, we performed cleavage under targets and release using nuclease (CUT&RUN) experiments in FBL-knockdown or SCR HUDEP-2 cells. We found that FBL was significantly enriched at transcription start sites (TSSs), implying that FBL may perform transcriptional regulatory functions (Fig. 4C). In particular, we found that FBL occupancy was highly enriched in the *HBG1/2* and *HBB* genes and in the locus control region (LCR), and accordingly, FBL enrichment in these regions was significantly reduced after FBL was knocked down (Fig. S7A). Notably, FBL occupancy was also markedly enriched in the BCL11A promoter and intronic enhancer regions (+55, +58, and +62) (*9, 13*) (Fig. S7B). These results indicated that FBL may also regulate the expression of BCL11A. We then assessed the occupancy of FBL-mediated H2AQ104me at the BCL11A gene via cleavage under targets and tagmentation (CUT&Tag) assays. We showed that FBL-mediated H2AQ104me was also markedly enriched in the BCL11A promoter region and the +55/+58 enhancer region (Fig. 4D). To examine whether FBL regulates chromatin accessibility, we carried out an assay for transposase-accessible chromatin with high-throughput sequencing (ATAC-seq) in FBL-depleted or SCR HUDEP-2 cells. A more closed chromatin state was shown at the BCL11A promoter and +55/+58 enhancer regions in FBL-depleted cells than in SCR cells (Fig. 4D), which is consistent with the transcriptome analysis. Real-time qPCR and western blot assays demonstrated that BCL11A mRNA and protein levels were significantly lower in FBL-depleted HUDEP-2 cells than in SCR cells (Fig. S7C), and KLF1, GATA1, and GATA2 mRNA levels remained unchanged (Fig. S7D). Indeed, the occupancy of RNA Pol II was reduced at the BCL11A promoter and +55/+58 enhancer regions after FBL disruption (Fig. S7E and F), reflecting diminished transcriptional activity. To examine the contribution of BCL11A to the induction of γ-globin upon FBL knockdown, we restored the expression of BCL11A in FBL-depleted HUDEP-2 cells by viral transduction of HA-tagged BCL11A. We found that exogenous HA-BCL11A overexpression partially reversed the derepression of γ-globin in FBL-depleted cells (Fig. S7G). Notably, the level of the histone marker H2AQ104me was not affected by HA-BCL11A (Fig. S7G). Furthermore, we knocked down BCL expression in FBL-depleted cells. We found that FBL/BCL11A double knockdown (DKD) led to a much greater increase in γ-globin mRNA and protein levels than FBL or BCL11A knockdown (Fig. 4E and F; Fig. S7H). Consistently, the percentage of HbF-expressing cells in the FBL/BCL11A double-knockdown group was nearly double that in the FBL- or BCL11A-knockdown groups (Fig. 4G). Taken together, these results indicate that FBL promotes BCL11A gene expression in erythroid cells.

### Repression of γ-globin expression by FBL is dependent on histone methyltransferase activity

FBL is a nucleolar protein that methylates histone H2A at glutamine 104 (H2AQ104me) (*25*). Since CUT&RUN assays have shown that FBL binds to HBG promoters, and we subsequently examined FBL-mediated H2AQ104me enrichment at the β-globin genes in HUDEP-2 cells via CUT&Tag assays. We showed that H2AQ104me was markedly enriched in the *HBG1/2* and *HBB* genes and in the LCR (Fig. 5A), which agrees with the FBL occupancy of the β-globin genes. Accordingly, FBL depletion led to a reduction in H2AQ104me in these regions (Fig. 5A). Subsequently, we tested chromatin accessibility across the β-globin locus by ATAC-seq. We found a significant reduction in chromatin accessibility at the *HBB* and LCR in FBL-depleted cells compared to SCR cells. Intriguingly, we found a significant increase in chromatin accessibility at the *HBG1* promoter in FBL-depleted cells compared to SCR cells (Fig. 5A). These results suggest that a reduction in H2AQ104me enrichment at the β-globin locus induces a switch from *HBB* to *HBG* expression.

**Fig. 5.**
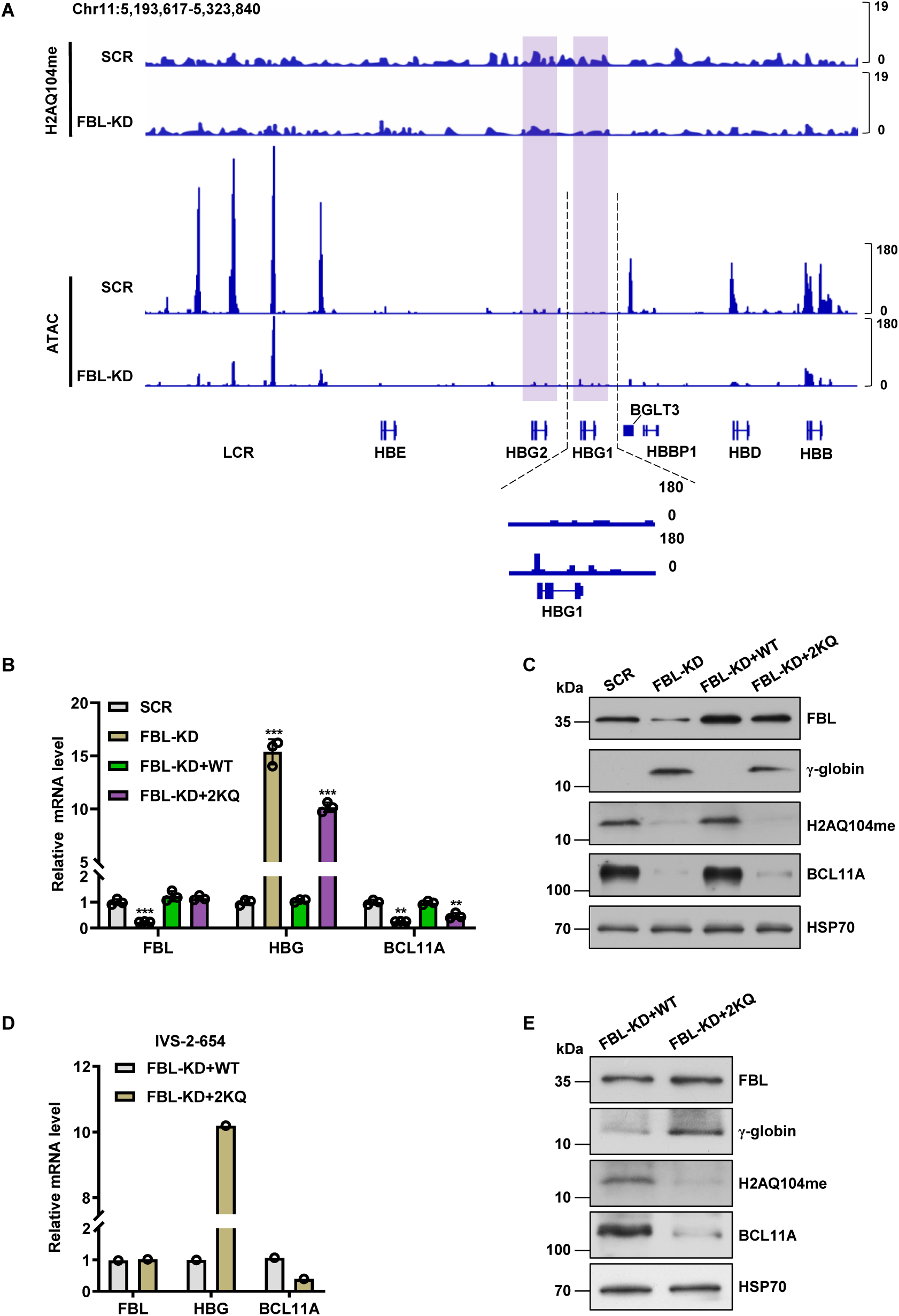
Repression of γ-globin expression by FBL is dependent on histone methyltransferase activity. **(A)** H2AQ104me enrichment analyzed by CUT&Tag and open chromatin regions analyzed by ATAC-seq at the β-globin locus in SCR or FBL-KD HUDEP-2 cells. **(B)** Real-time qPCR analysis of *FBL*, *HBG,* and *BCL11A* mRNA expression in SCR, FBL-KD, FBL-KD+WT, and FBL+2KQ HUDEP-2 cells. The results were normalized to GAPDH mRNA and are shown as the mean ± SD from three independent experiments. \*\**P* < 0.01; \*\*\**P* < 0.001. **(C)** Western blot of FBL, γ-globin, H2AQ104me, and BCL11A in SCR, FBL-KD, FBL-KD+WT, or FBL+2KQ HUDEP-2 cells. **(D)** Real-time qPCR showing the expression of *FBL*, *HBG,* and *BCL11A* mRNAs in FBL-KD+WT and FBL+2KQ CD34^+^ HSPCs from β-thalassemia patients. The results were normalized to GAPDH mRNA and are shown as the mean ± SD from three independent experiments. \*\**P* < 0.01; \*\*\**P* < 0.001. **(E)** Western blot of FBL, γ-globin, H2AQ104me, and BCL11A in FBL-KD+WT and FBL+2KQ CD34^+^ HSPCs from β-thalassemia patients.

Among nucleolar proteins, FBL is an essential protein that mediates H2AQ104me as well as 2’-O-methylation of rRNA (*25, 35–37*). Studies have shown that the replacement of lysine residues (K102, K121, K205, and K206) in FBL with glutamine (4KQ) disrupts FBL-mediated H2AQ104me without affecting the 2’-O-methylation of rRNA (*38*). Lysine residues K205 and K206 are located in the methyltransferase domain, whereas K102 and K121 of FBL are located in the spacer 1 domain (*26, 38*). To confirm whether the regulation of γ-globin expression by FBL was histone methyltransferase activity dependent, we constructed a mutant form of FBL (FBL-2KQ) in which two lysines at positions 205 and 206 were mutated to glutamines (K205Q and K206Q) (*26, 38*). Western blot analysis confirmed the inability of FBL-2KQ to methylate H2AQ104 (Fig. 5C). As expected, transfection of wild-type FBL into FBL-knockdown cells restored H2AQ104me levels (Fig. 5B and C), but, transfection of FBL-2KQ into FBL-knockdown cells did not (Fig. 5C). We observed that BCL11A expression was restored in FBL-depleted cells by wild-type FBL but not by the FBL-2KQ mutant (Fig. 5B and C). Accordingly, γ-globin expression was repressed by wild-type FBL in FBL-depleted cells but derepressed by the FBL-2KQ mutant.

Next, we tested whether the enzymatically inactive FBL mutant could function in HSPCs from β-thalassemia patients with the IVS-2-654 mutation. Compared with wild-type FBL (FBL-WT), FBL-2KQ overexpression significantly induced γ-globin expression in FBL-knockdown cells (Fig. 5D and E). Accordingly, we detected lower levels of H2AQ104me and BCL11A in FBL-knockdown cells reconstituted with FBL-2KQ than in those reconstituted with FBL-WT (Fig. 5E). These results indicate that the repression of γ-globin expression by FBL is dependent on histone methyltransferase activity.

## Discussion

In this study, we identified FBL as a novel γ-globin gene repressor. We found a dual role for FBL, which activates BCL11A expression and directly represses γ-globin expression in erythroid cells. Our current findings add another key γ-globin silencer harboring effects comparable to those of BCL11A, LRF, and NFI factors (*5, 8, 9*). Our work supports a model in which complete γ-globin silencing requires the coregulation of these factors.

The interaction of transcription factors with epigenetic regulators controls gene expression. Previously, we showed that LYAR interacts with PRMT5 to repress γ-globin expression (*19–22*). The common rSNP rs368698783 A-allele has been found to attenuate LYAR and the epigenetic regulators DNMT3A and PRMT5 from the HBG promoter, resulting in elevated HbF levels in individuals with β-thalassemia (*22, 39*). It would be interesting to reevaluate the contribution of FBL in this context. Given that FBL binds well to the β-globin gene locus, we cannot exclude the possibility that FBL may cooperate with other transcription factors or regulators to modulate the expression of β-globin genes.

While FBL depletion led to a reduced enrichment of H2AQ104me at the *BCL11A* enhancer regions (+55 and +58) and the promoter region, a more closed chromatin at these regions was observed in FBL knockdown cells. In fact, FBL enrichment correlated well with RNA polymerase II enrichment on the *BCL11A* gene. Therefore, the H2AQ104me mark seems to be an active gene marker, although the detailed mechanism remains to be determined. A similar scenario was observed for the LCR, *HBD*, and *HBB* genes in HUDEP cells. Although FBL depletion led to a reduced enrichment of FBL at the LCR, *HBD*, and *HBB* and a more closed chromatin at these loci, a more open chromatin at *HBG* (particularly *HBG1*) was observed even if a reduction in H2AQ104me enrichment occurred at the *HBG* promoter. This pattern supports the gene competition model of hemoglobin switching at this stage (*40, 41*). In this context, H2AQ104me may weaken the interaction between H2A and the FACT complex, which may interact with RNA polymerase II (*25*). Nevertheless, the specific regulation of gene repression associated with FBL-mediated H2AQ104me warrants further investigation.

The βIVS-2-654 mutation is one of the most common mutations in the Chinese population accounting for approximately 20% of all β-thalassemia cases in southern China. The correction of the phenotype of IVS-2-654 β-thalassemic mice and CD34^+^ HSPCs from β-thalassemia patients upon depletion of FBL provides both in vitro and in vivo evidence and proof of principle for therapeutic targeting of FBL in β-thalassemia patients. We anticipate that our findings should apply similarly to SCD patients. Our study revealed that interference with FBL expression or methyltransferase activity may be pursued to help treat patients with β-thalassemia and SCD.

## Acknowledgments

We thank the members of the Zhao Laboratory for their helpful discussions. This work was supported by the National Natural Science Foundation of China (NSFC #32270619 and #31970615), Fundamental Research Fund of the State Key Laboratory of Pharmaceutical Biotechnology (#LNSN-202403).

