## Supplementary material for "Fibrillarin modulates fetal hemoglobin silencing": Table S1

[1]The State Key Laboratory of Pharmaceutical Biotechnology, Department of Hematology, the Affiliated Drum Tower Hospital of Nanjing University Medical School, China-Australia Institute of Translational Medicine, School of Life Sciences, Nanjing University, Nanjing, China; [2]Department of Hematology, Jinling Hospital, School of Medicine, Nanjing University, Nanjing, China; [3]The Walter and Eliza Hall Institute of Medical Research, Department of Medical Biology, University of Melbourne, Melbourne, VIC, Australia.

**This file includes:**

Materials and methods
Figs. S1 to S7
Tables S1 to S2
References

#### Materials and methods

##### Cell culture and erythroid maturation

HEK293T cells were cultured in DMEM containing 10% FBS and 1% penicillin–streptomycin. K562 cells were cultured in RPMI-1640 medium supplemented with 10% FBS and 1% penicillin–streptomycin.

HUDEP-2 cells were cultured in StemSpan SFEM II (StemCell Technologies, 09655) medium supplemented with 50 ng/ml recombinant human stem cell factor (SCF) (MedChemExpress, HY-P7056), 3 IU/ml erythropoietin (R&D Systems), 1 μg/ml doxycycline (MedChemExpress), and $10^{-6}$ M dexamethasone (MedChemExpress).

$CD34^+$ HSPCs were obtained from adult human peripheral blood or bone marrow and cultured in StemSpan SFEM II containing 1×CC100 cytokine mix (StemCell Technologies, 02690) for 6 days. For erythroid differentiation, phase I Iscove's modified Dulbecco's medium (IMDM, Thermo Fisher, 12440053) supplemented with 3 IU/ml erythropoietin, 1 ng/ml interleukin-3 (StemCell Technologies), 10 μg/ml insulin (Sigma, I9278), 200 μg/ml holo-transferrin (Sigma, T0665), 5% human AB plasma (SeraCare), 100 ng/ml human SCF and 2 IU/ml heparin (MedChemExpress, HY-17567A) was used. In phase II, interleukin-3 was omitted. 8 days in Phase I and 4 days in Phase II. Anti-CD235a (BD, 555570) and anti-CD71 (BD, 551374) antibodies were used to analyze erythroid maturation by flow cytometry. Cells were authenticated by short tandem repeat (STR) profiling and were negative for mycoplasma contamination.

##### Cell growth assay

$CD34^+$ HSPCs were cultured in 96-well plates at 2000 cells per well (200 μl). Ten microliters of Cell Counting Kit-8 (CCK-8, ApexBio, K1018) solution was added to the wells. After incubation for 2 h, the absorbance at 450 nm was measured.

**Western blot**

After being washed with PBS, the cells were lysed in cell lysis buffer (Beyotime, P0013) and mixed with 5×protein loading buffer. Proteins were separated by SDS–PAGE, transferred to PVDF membranes (Roche, 03010040001), incubated in blocking solution, and then incubated with each primary antibody overnight. After being washed with TBS-T (Tris-buffered saline with 0.1% Tween-20 buffer), the membranes were incubated with secondary antibodies for 2 h at room temperature. Primary antibodies against FBL (Abcam, ab166630), BCL11A (Abcam, ab191401), H2AQ104me (Active motif, 61605), γ-globin (Abcam, ab137096), and LYAR (ABclonal, A17724) were used.

**Mass spectrometry and protein interaction assays**

SDS–PAGE gels were used to isolate the FLAG antibody immunoprecipitate of LYAR-FLAG-overexpressing K562 cells, which were subsequently stained with Coomassie brilliant blue. The protein bands were obtained and analyzed by LC–MS/MS. Coimmunoprecipitation and glutathione S-transferase (GST) pulldown assays were performed according to previous methods (*1*).

**Lentivirus or retrovirus infection and shRNA lentiviral transduction**

FBL, LYAR, and BCL11A were knocked down by shRNA. The sequences were cloned and inserted into the pLL3.7 lentiviral vector. The sequences were as follows:

FBL-KD1 (human): GATTTCGGAAGGAGATGACAA

FBL-KD2 (human): CCTTGAGCCATATGAAAGAGA

LYAR-KD (human): CCTGGTCATCTTTAACAAG

human BCL11A-KD: ACGCACAGAACACTCATGGATT

The rescue of BCL11A was achieved by cloning BCL11A cDNA into pSDM101. To overexpress FBL, human FBL cDNA was cloned and inserted into a retroviral vector plasmid (MSCV-HA-IRES-GFP). 293T cells were infected with lentivirus or retrovirus. Briefly, the shRNA lentiviral vector, psPAX2 and pMD2.G packaging plasmids were mixed in DMEM supplemented with DNAfectin™ Plus Transfection Reagent (Applied Biological Materials, G2500) and then cotransfected into 293T cells. After 48 h of transfection, the supernatants containing the virus particles were used to infect HUDEP-2 cells.

**Wright-Giemsa Staining**

The cells were collected and washed with PBS, and then the cells were spun on slides by centrifugation. The cells were fixed in 70% ethanol for 10 min and stained with staining solution (Beyotime, C0131) for 45 min. After washing with deionized water, the slides were microscopically observed and imaged.

**HbF staining and flow cytometry analysis**

Briefly, cells were collected and washed with PBS, treated with 0.05% glutaraldehyde for 10 min, washed with PBS, and then permeabilized using 0.1% Triton X-100 for 3 min. Finally, the cells were washed, resuspended in PBS, and then incubated with an APC-conjugated HbF antibody (Thermo Fisher, MHFH05) in the dark for 15-20 min. The percentage of HbF-immunostained F-cells was detected by flow cytometry and analyzed via FlowJo.

**Hemoglobin analysis**

Two million cells were collected for hemoglobin analysis after differentiation. The cells were washed with PBS and lysed in 200 μl of Milli-Q $H_2O$. The supernatant containing the proteins was obtained by centrifugation, the cell debris was discarded, and the levels of different hemoglobins were analyzed by HPLC (Bio-Rad, D-10$^{TM}$).

**Real-time qPCR and ChIP analysis**

Briefly, one million cells were collected, and RNA was extracted after TRIzol (Thermo Fisher, 15596018CN) treatment. RNA (200-600 μg) was used to prepare cDNA with a HiScript cDNA kit (Vazyme). Real-time qPCR was performed via a QuantStudio 3 Real-Time PCR instrument (Thermo Fisher) using SYBR Green master mix (Thermo Fisher). The ChIP assay was performed according to a previous method (*1*). All primers used for real-time qPCR or ChIP assays are listed in Tables S1 and S2.

**RNA-seq**

To analyze the changes in genome-wide gene expression in HUDEP-2 cells after FBL knockdown, we constructed SCR and FBL-KD HUDEP-2 cell lines. Each group contained three repeated samples. The extraction of RNA was performed according to previous methods. RNA-Seq libraries were prepared according to the instructions of the TruSeq Stranded Total RNA Kit (Illumina). After quality testing, the library was sequenced on a Novaseq XPlus (Illumina) platform. DESeq2 (1.20.0) was used to perform differential expression analysis between the SCR and FBL-KD groups.

**CUT&RUN and CUT&Tag assay**

A total of $2\times10^4$ to $5\times10^5$ cells were used for the CUT&RUN assay. We generated libraries using a CUT&RUN Assay Kit (Vazyme). We carried out the assay according to the method published by the laboratory of Henikoff (2). Briefly, the treated cells were incubated with antibodies for 2 hours or overnight. The antibodies used were anti-FBL (Abcam, ab5821). DNA fragmentation was performed by the pAG-MNase Enzyme (Cell Signaling). The target DNA fragments were sorted from the products and sequenced on the NovaSeq XPlus platform (Illumina). For the CUT&Tag assay, we generated libraries using a CUT&Tag Assay Kit (Vazyme). Anti-H2AQ104me (Active Motif, 61605) was used as the primary antibody, and the secondary antibody was obtained from Abcam. Finally, the libraries were sequenced, and bigwig files were generated.

**ATAC-seq**

A total of $5\times10^4$ to $9\times10^5$ cells were centrifuged at 2300 rpm for 5 min at 4°C, and the supernatant was discarded. Each sample was resuspended in precooled TW buffer (tagment DNA and wash buffer, Vazyme) and centrifuged at 2,300 rpm for 5 min at 4°C, after which the supernatant was discarded. The cells were resuspended in lysis buffer, placed on ice for 5 min, and centrifuged at 2,300 rpm for 10 min at 4°C, after which the supernatant was discarded. Each sample was resuspended in precooled fragmentation mix at 37°C for 30 min. The cells were subsequently resuspended in stop buffer (Vazyme), after which ATAC-DNA-Extract Beads were added to extract DNA from the sample. The obtained DNA was subjected to PCR according to the instructions of the Hyperactive ATAC-Seq Library Prep Kit (Vazyme). Finally, ATAC-DNA-Clean Beads were used to sort target fragments from the PCR products. Libraries were sequenced, and bigwig files were generated.

**Animal studies**

β-YAC and Mx1-*Cre* mice have been previously characterized (*3,4,5*). Mx1-*Cre* and *Fbl*$^{fl/-}$ mice were obtained from Shanghai Model Organisms in China, and the experiments were approved by the Animal Ethical and Welfare Committee of Nanjing University. The sample size was chosen with adequate power based on the literature and our previous experience; for each experiment, sample size is indicated in the figure legend. Prior to the experiment, the mice were randomly assigned to different treatment groups. We obtained *Fbl*$^{fl/fl}$::Mx1-*Cre*$^{+}$::β-YAC or *Fbl*$^{fl/fl}$::Mx1-*Cre*$^{-}$::β-YAC mice by crossbreeding. Poly (I:C), an inducer, was dissolved in PBS at a concentration of 2 mg/ml, and mice were intraperitoneally injected 4 times with 25 mg/kg poly(I:C) administered every other day. After induction, the mice were allowed to recover for one or two weeks in a stable environment. Then, we collected peripheral blood from the mice for complete blood count analysis and bone marrow for western blot analysis, real-time qPCR, and flow cytometry. Anti-mouse Ter119 (BioLegend, 116207) and anti-mouse CD71 (BioLegend, 113805) antibodies were used to analyze erythroid maturation by flow cytometry.

β$^{IVS-2-654}$-thalassaemic mice were obtained from Jackson Laboratory. The AAV was packaged in AAV-293 cells. β-Thalassemia mice (8 weeks old) received an intramedullary injection of 5 μl of AAV (Shanghai Genechem Co., Ltd.). After 8 weeks, we collected 200 μl of peripheral blood from the mice for complete blood count analysis using an XT-2000i (Sysmex) and 3-5 μl of peripheral blood for blood smears. Blood smears were stained with Wright and Giemsa (Beyotime, C0131) and analyzed by microscopy. The bone marrow was collected for western blot analysis and real-time qPCR. Next, we obtained spleen images and recorded weights, and spleens were analyzed by hematoxylin-eosin (H&E) staining.

**Statistical analysis**

All the data are presented as means ± SDs unless otherwise indicated. GraphPad Prism software (version 7.0) was used to assess significance of differences. Comparisons between two groups were performed using an unpaired two-tailed Student's t test. Multiple comparisons were performed using one-way analysis of variance (ANOVA); $P < 0.05$ was considered significant. In the graphics, one, two, and three asterisks indicate $P < 0.05$, $P < 0.01$, and $P < 0.001$, respectively.

**Data availability**

All relevant data are available within the article and supplementary files or are available from the authors upon request. The raw mass spectrometry proteomics data were deposited in the ProteomeXchange consortium under accession number PXD053272. The RNA-Seq, ATAC-Seq, CUT&RUN and CUT&Tag data generated in this study have been deposited in the Gene Expression Omnibus (GSE270538, GSE270535, GSE270536, and GSE270537).

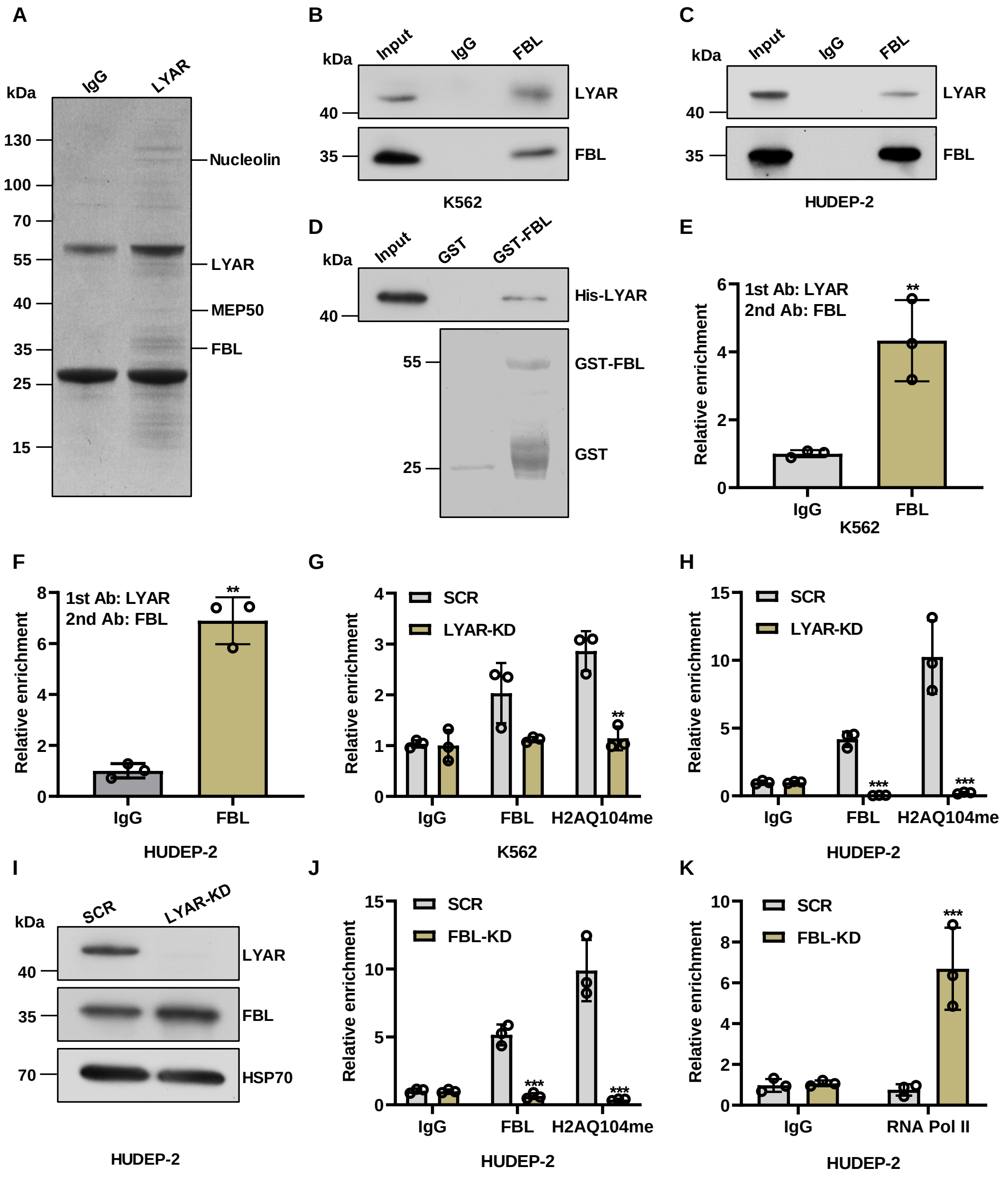


#### Figure S1

**Fig. S1: FBL interacts with LYAR. (A)** Coomassie brilliant blue (CBB)-stained gel of proteins separated by SDS–PAGE after anti-FLAG M2 affinity gel or IgG (control) immunoprecipitation from K562 cells stably overexpressing FLAG epitope-tagged LYAR before analysis by mass spectrometry. The bands corresponding to FBL, LYAR, MEP50, and Nucleolin are shown. **(B, C)** Endogenous FBL was coimmunoprecipitated with LYAR in K562 (**B**) and HUDEP-2 cells (**C**). Western blot was probed with anti-

LYAR antibodies. IgG served as a negative control. **(D)** GST pull-down assay (upper panel). Prokaryotic GST and GST-FBL fusion proteins preabsorbed to glutathione-Sepharose beads were incubated with purified prokaryotic His-LYAR fusion protein. Specifically, bound protein was eluted from the washed beads and visualized by Western blot analysis with an anti-LYAR antibody after SDS–PAGE. Coomassie-stained SDS–PAGE gel showing purified prokaryotic GST and GST–FBL fusion proteins (lower panel). **(E, F)** ChIP-reChIP (anti-LYAR antibody ChIP followed by anti-FBL antibody ChIP) analysis of LYAR and FBL on the γ-globin promoter in K562 (**E**) or HUDEP-2 cells (**F**). Normal IgG served as the control. The results are shown as the mean ± SD from three independent experiments. $^{**}P < 0.01$. **(G, H)** ChIP analysis of the enrichment of FBL or H2AQ104me on the γ-globin promoter in K562 (**G**) or HUDEP-2 cells (**H**). IgG served as a negative control. The results are shown as the mean ± SD from three independent experiments. $^{**}P < 0.01$, $^{***}P < 0.001$. **(I)** Western blot analysis of LYAR and FBL expression in LYAR-KD or SCR HUDEP-2 cells. **(J)** ChIP analysis of the enrichment of FBL or RNA Pol II on the γ-globin promoter in FBL-KD or SCR HUDEP-2 cells. IgG served as a negative control. The results are shown as the mean ± SD from three independent experiments. $^{***}P < 0.001$. (**K**) ChIP analysis of the enrichment of RNA Pol II on the γ-globin promoter in FBL-KD or SCR HUDEP-2 cells. IgG served as a negative control. The results are shown as the mean ± SD from three independent experiments. $^{***}P < 0.001$.

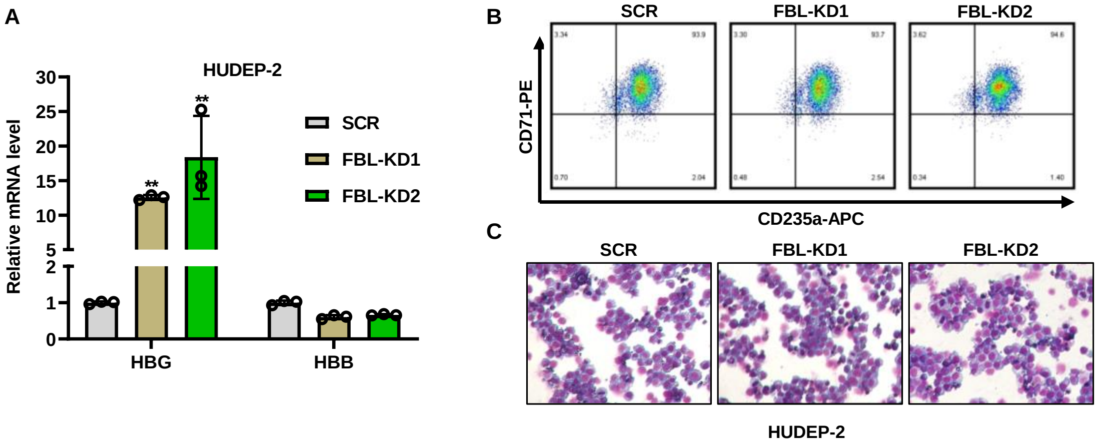


### Figure S2

**Fig. S2: FBL disruption induces HBG expression in HUDEP-2 cells. (A)** Real-time qPCR showing the expression of *HBG* and *HBB* in the indicated HUDEP-2 cells. The results are shown as the mean ± SD from three independent experiments. $^{**}P < 0.01$. **(B)** CD71 and CD235a were analyzed by flow cytometry in SCR or FBL-KD HUDEP-2 cells. **(C)** Wright–Giemsa staining of SCR or FBL-KD HUDEP-2 cells.

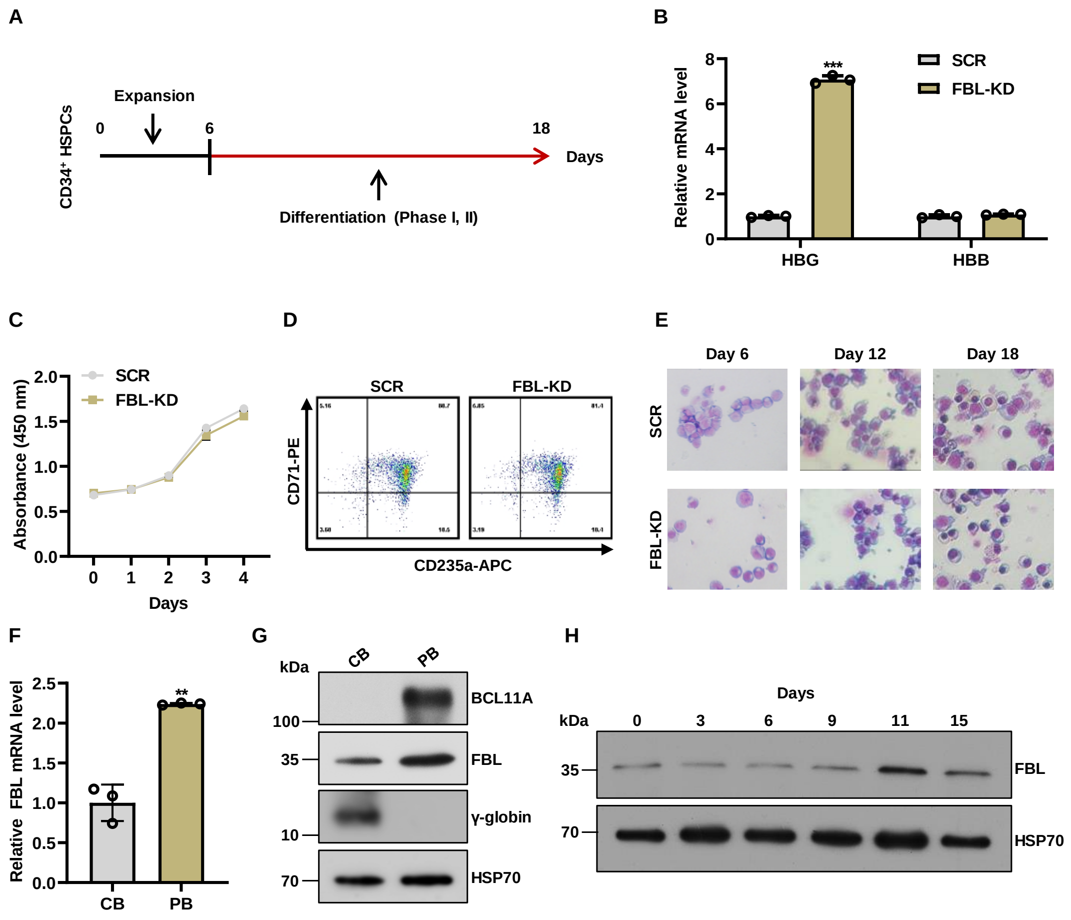


### Figure S3

**Fig. S3: FBL disruption induces HBG expression in CD34⁺ HSPCs. (A)** Schematic diagram of the CD34⁺ HSPC culture and differentiation system. **(B)** Real-time qPCR showing the expression of *HBG* and *HBB* in the indicated CD34⁺ HSPCs. The results are shown as the mean ± SD from three independent experiments. ***$P < 0.001$. **(C)** Cell expansion curves of SCR and FBL-KD CD34⁺ HSPCs. The results are shown as the mean ± SD from three independent experiments. **(D)** CD71 and CD235a were analyzed by flow cytometry in SCR or FBL-KD CD34⁺ HSPCs. **(E)** Wright–Giemsa staining of SCR or FBL-KD CD34⁺ HSPCs at different time points. **(F)** Real-time qPCR analysis of FBL expression in cord blood (CB, newborn) and peripheral blood

(PB, adult) erythroblasts. The results were normalized to GAPDH mRNA and are shown as the mean ± SD from three independent experiments. ***P* < 0.01. **(G)** Western blot analysis of BCL11A, FBL and γ-globin in CBs and PBs. HSP70 served as a loading control. **(H)** Western blot analysis of FBL in $CD34^+$ HSPCs on the indicated days. HSP70 served as a loading control.

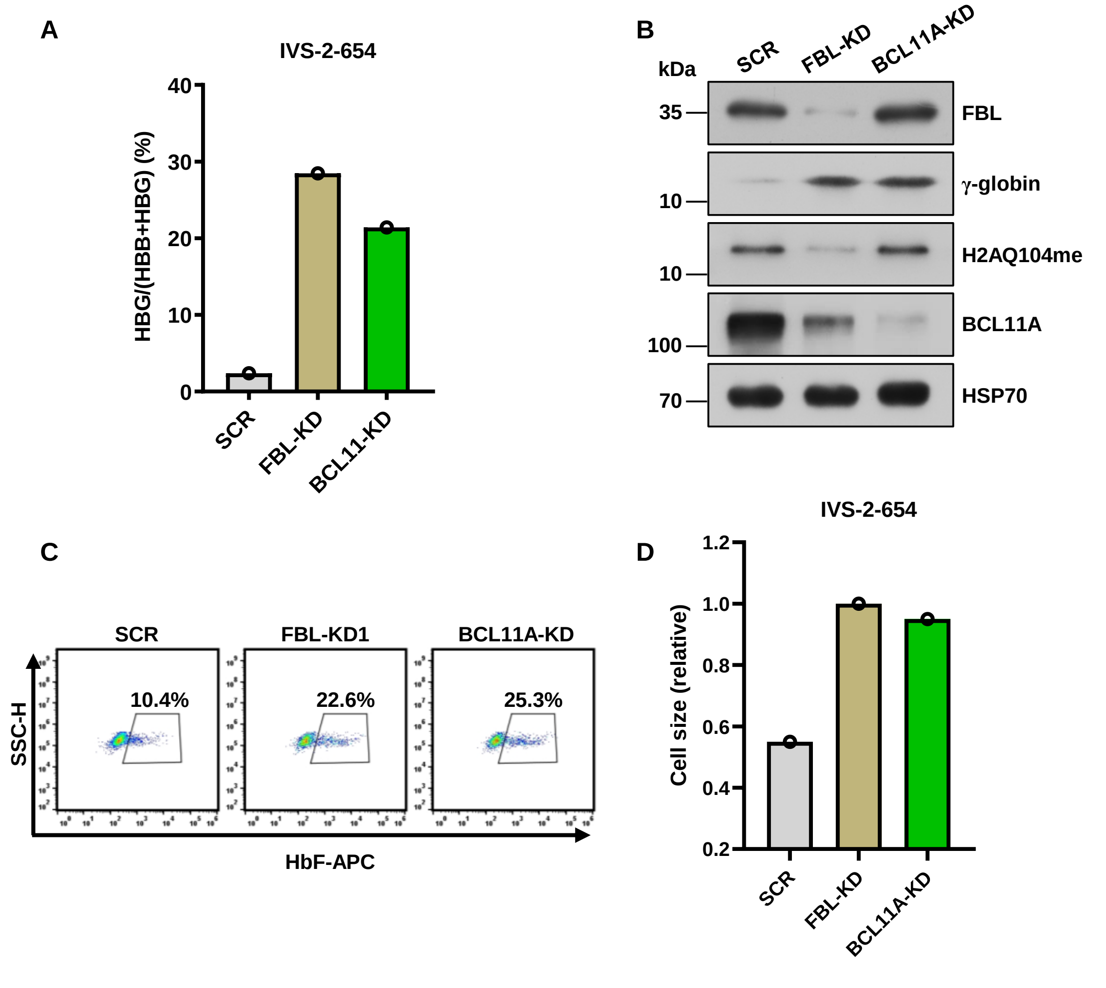


### Figure S4

**Fig. S4: FBL disruption induces γ-globin expression in primary human erythroid progenitor cells from β-thalassemia patients. (A)** Real-time qPCR showing the ratio of HBG/(HBG + HBB) in $CD34^+$ HSPCs from IVS-2-654 β-thalassemia patients (n = 1 donor). **(B)** Western blot analysis of FBL, γ-globin, H2AQ104me, and BCL11A in $CD34^+$ HSPCs from IVS-2-654 β-thalassemia patients (n = 1 donor). **(C)** The percentage of HbF-immunostained F-cells among the indicated cells was detected by flow cytometry (n = 1 donor). **(D)** Cell size was measured by relative forward scatter intensity for erythroid cells differentiated from IVS-2-654 β-thalassemia patients (n = 1 donor).

A

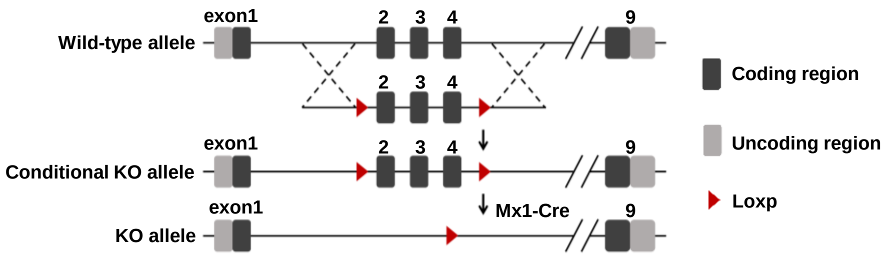


B

| Mice | WBC ×10⁹/L | RBC ×10¹²/L | HGB g/L | HCT % | MCV fL | MCH pg | MCHC g/L | RDW % | PLT ×10⁹/L | MPV fL |
|---|---|---|---|---|---|---|---|---|---|---|
| Control | 5.69 | 8.81 | 125 | 49.3 | 56.0 | 14.1 | 253 | 32.5 | 935 | 6.3 |
| $Fbl^{-/-}$ | 4.17 | 8.89 | 124 | 46.6 | 52.5 | 14 | 266 | 33 | 620 | 6 |

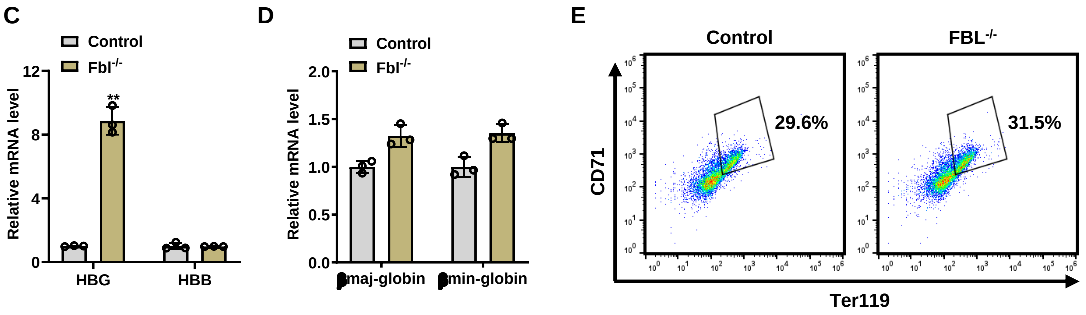


### Figure S5

**Fig. S5: Fbl depletion in adult β-YAC mice. (A)** Schematic diagram of Mx1-Cre-mediated excision of exons 2 to 4 (flanked by loxP sites) in the Fbl locus to produce the Fbl mutant allele. **(B)** Hematologic parameters of control (n=3) and *Fbl*$^{-/-}$ knockout (n=3) mice. The results are shown as the mean ± SD. WBC, white blood cell; RBC, red blood cell; HGB, hemoglobin; HCT, hematocrit; MCV, mean corpuscular volume; MCH, mean corpuscular hemoglobin; MCHC, mean corpuscular hemoglobin concentration; RDW, red blood cell distribution width; PLT, platelet; MPV, mean platelet volume. **(C)** Real-time qPCR showing the expression of *HBG* and *HBB* mRNAs in bone marrow from control or *Fbl*$^{-/-}$ knockout mice. The results were normalized to

GAPDH mRNA and are shown as the mean ± SD from three independent experiments. $^{**}P < 0.01$. **(D)** Real-time qPCR showing the expression of adult β-globulin (βmajor and βminor) mRNAs in bone marrow from control or *Fbl*$^{-/-}$ knockout mice. The results were normalized to GAPDH mRNA and are shown as the mean ± SD from three independent experiments. $^{**}P < 0.01$. **(E)** The erythroid markers CD71 and Ter119 were measured in bone marrow cells from the indicated mice by flow cytometry.

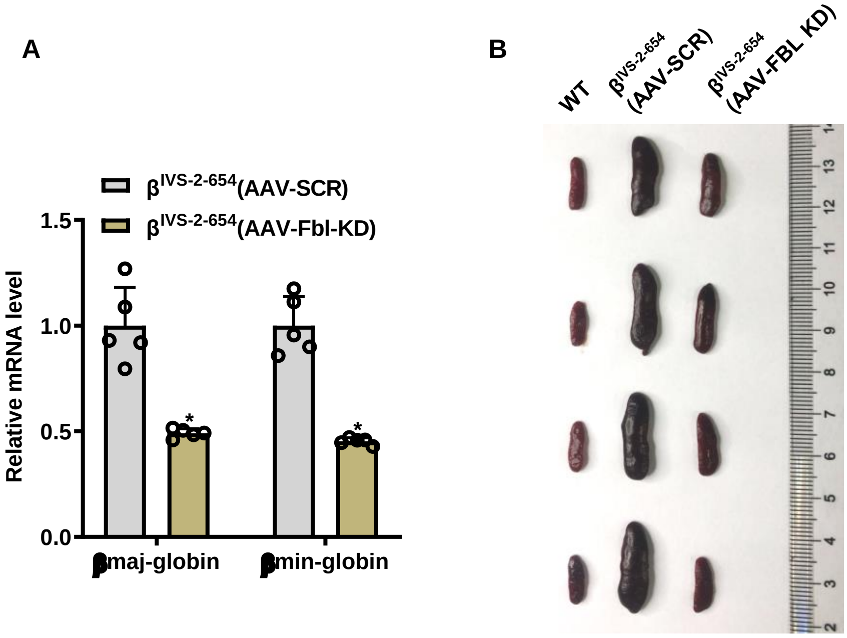


### Figure S6

**Fig. S6: Depletion of Fbl corrects anemia in a mouse model of beta-thalassemia.** **(A)** Real-time qPCR showing the expression of mouse adult β-globins (βmajor and βminor) in AAV-SCR- or AAV-FBL-KD-treated thalassemic mice. The results were normalized to β-actin mRNA and are shown as the mean ± SD from five independent experiments. *$P < 0.05$. **(B)** Spleen images of WT, AAV-SCR-, or AAV-FBL-KD-treated thalassemic mice.

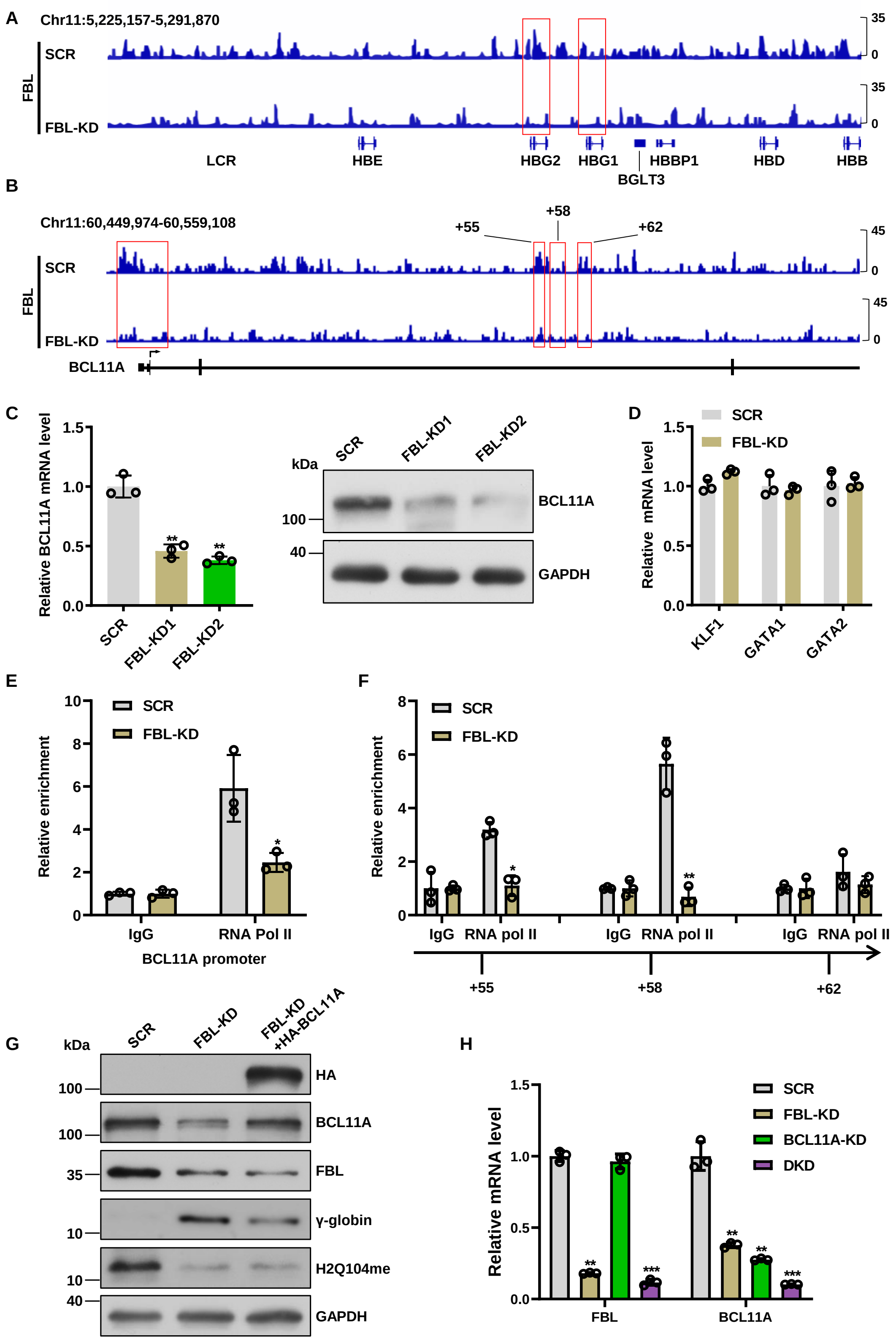


### Figure S7

**Fig. S7: FBL supports the expression of BCL11A. (A and B)** Enrichment of FBL analyzed by CUT&RUN at the β-globin (**A**) and BCL11A (**B**) loci in SCR or FBL-KD

HUDEP-2 cells. **(C)** Real-time qPCR showing the expression of FBL mRNA in SCR, FBL-KD1, or FBL-KD2 HUDEP-2 cells (left panel). The results are shown as the mean ± SD from three independent experiments. **$P < 0.01$. Western blot analysis of FBL protein in the indicated cells. GAPDH served as a loading control (right panel). **(D)** Real-time qPCR showing the mRNA levels of KLF1, GATA1, and GATA2 in SCR or FBL-KD HUDEP-2 cells. The results are shown as the mean ± SD from three independent experiments. **(E)** The enrichment of RNA Pol II at the BCL11A promoter in SCR or FBL-KD HUDEP-2 cells was analyzed by ChIP assay. The results are shown as the mean ± SD from three independent experiments. *$P < 0.05$. **(F)** The enrichment of RNA Pol II at the BCL11A +55, +58, and +62 enhancer regions in SCR or FBL-KD HUDEP-2 cells was analyzed by ChIP. The results are shown as the mean ± SD from three independent experiments. *$P < 0.05$; **$P < 0.01$. **(G)** Western blot analysis of BCL11A, FBL, γ-globin, and H2AQ104me in SCR, FBL-KD, or FBL-KD+HA-BCL11A HUDEP-2 cells. GAPDH served as a loading control. **(H)** Real-time qPCR analysis of *FBL* and *BCL11A* mRNA expression in the indicated cells. The results are shown as the mean ± SD from three independent experiments. **$P < 0.01$; ***$P < 0.001$.

**Table S1.** Primers used for real-time qPCR

| Gene name | Species | Forward primer (5′-3′) | Reverse primer (5′-3′) |
|---|---|---|---|
| FBL | Human | CATTCTTCCCCGACTGGTTTC | GTCGAGGCGGAGGCTTTAG |
| BCL11A | Human | CGCCAGAGGATGACGATTGTT | CCAGGCGTGGGGATTAGAG |
| GAPDH | Human | ACCCAGAAGACTGTGGATGG | TTCAGCTCAGGGATGACCTT |
| GATA1 | Human | AGTTTGTGGATCCTGCTCTG | GCAATGGGTACACCTGAAAG |
| GATA2 | Human | GCAACCCCTACTATGCCAACC | CAGTGGCGTCTTGGAGAAG |
| KLF1 | Human | GGTTGCGGCAAGAGCTACA | GTCAGAGCGCGAAAAAGCAC |
| HBG | Human | AATGTGGAAGATGCTGGAGGAGAA | CTTCTTGCCATGTGCCTTGACTT |
| HBB | Human | CTGAGGAGAAGTCTGCCGTTA | AGCATCAGGAGTGGACAGAT |
| εy-globin | Mouse | TGGCCTGTGGAGTAAGGTCAA | GAAGCAGAGGACAAGTTCCCA |
| βh1-globin | Mouse | GAAACCCCCGGATTAGAGCC | GAGCAAAGGTCTCCTTGAGGT |
| βmaj-globin | Mouse | GGGTAATGCCAAAGTGAAGGC | GGCCCAGCACAATCACGATCAT |
| βmin-globin | Mouse | TCTGCTGTCTCTTGCCTGTG | CCTTTTTGCCATGGGCCTTC |
| β-actin | Mouse | GGCTGTATTCCCCTCCATCG | CCAGTTGGTAACAATGCCATGT |

**Table S2.** Primers used for ChIP-qPCR

| Gene name | Forward primer (5′-3′) | Reverse primer (5′-3′) |
|---|---|---|
| HBG (**-260~-1**) | GAATCGGAACAAGGCAAAGG | GTGGAACTGCTGAAGGGTG |
| BCL11A-55 | GCACCTGCATTTGTTTTTCA | GGGTCAGATCACCTCTGCTC |
| BCL11A-58 | TGGACTTTGCACTGGAATCA | GATGGCTGAAAAGCGATACA |
| BCL11A-62 | TTTCAACCATGGTCATCTGC | CCCTCTGGCATCAAAATGAG |
